# Inhibition of Acyl-CoA: Cholesterol Acyltransferase 1 Promotes Shedding of Soluble TREM2 and LRP1-Dependent Phagocytosis of Amyloid β Protein in Microglia

**DOI:** 10.1101/2025.06.12.659424

**Authors:** Moriah J. Hovde, Anna Maaser-Hecker, Rudolph E. Tanzi

## Abstract

Lipid regulation plays a major role in the pathogenesis of Alzheimer’s disease (AD). In AD patients and transgenic mice, microglia exhibit increased expression of *SOAT1*, encoding Acyl-CoA: Cholesterol Acyltransferase 1 (ACAT1), which catalyzes the production of cholesteryl esters in lipid droplets. ACAT1 Inhibition has been reported to reduce β-amyloid pathology. However, the molecular mechanism underlying this effect remains unknown. Here, we show ACAT1 inhibition in mouse and human iPSC-derived microglia upregulates LRP1 levels and shedding of soluble TREM2 (sTREM2) owing to enhanced cleavage of TREM2 by ADAM10/17. Knocking out TREM2 or preventing sTREM2 release abrogated the ability of ACAT1 inhibition to enhance microglial uptake of amyloid β (Aβ). This could be rescued by the addition of recombinant sTREM2, but only in the presence of LRP1. Collectively, these data indicate ACAT1 inhibition increases microglial uptake of Aβ in a sTREM2- and LRP1-dependent manner suggesting new avenues for treating and preventing AD.

## Introduction

The amyloid β protein (Aβ) plays a central role in the etiology and pathogenesis of Alzheimer’s disease (AD), the most common form of dementia in the elderly^1^. Aβ accumulates as oligomers and plaques triggering a neurodegenerative process that leads to cognitive decline^2^. In addition to amyloid plaques, neurofibrillary tangles (NFT) and neuroinflammation, Alois Alzheimer reported the accumulation of lipid droplets in the glial cells of AD patients^3^, a finding largely overlooked for many years. A resurgence of interest in the many roles of lipid metabolism has inspired multiple studies investigating the role of lipid and cholesterol metabolism in AD. Regulation of cholesterol homeostasis occurs primarily in the endoplasmic reticulum (ER)^4^, a cholesterol poor-organelle which contains many of the enzymes responsible for both cholesterol synthesis and storage^5^. Cholesterol storage is predominantly driven by the resident ER protein Acyl-CoA: Cholesterol Acyltransferase 1 (ACAT1)^6^ which catalyzes esterification of membrane-associated cholesterol with a fatty acyl from acyl-CoA to produce cholesteryl esters (CE)^7,8^, which are then targeted to cytosolic lipid droplets^9,10^.

Lipid droplets are primarily composed of two classes of lipids: cholesteryl esters (CEs) and triacylglycerols (TAGs) which make up the neutral lipid core of the mono layered droplet. Brain samples from late-onset AD (LOAD) patients exhibit a 1.8-fold increase in CE levels versus controls, in vulnerable brain regions^11^. Additionally, multiple AD mouse models have shown 3- to 11-fold higher CE levels in the brain compared to control^11,12^. Finally, microglia isolated from vulnerable brain regions in various neurogenerative diseases, including LOAD, contain elevated levels of ACAT1 (*SOAT1*) mRNA^13^. These data have positioned ACAT1 as an attractive therapeutic target for AD.

ACAT1 inhibitors, originally developed for managing plasma cholesterol have been shown to reduce Aβ-induced neurotoxicity in both *in vitro* and *in vivo* mouse models (reviewed in ^14,15^). Previous *in vivo* work from our lab shows ACAT1 inhibition reduces Aβ production and promotes plaque clearance in AD mouse models^16^. Others have shown ACAT1 inhibition reduces AD pathology by promoting microglial-mediated clearance of Aβ^17–19^. Furthermore, reducing CE levels via ACAT1 inhibition in microglia cells that have been overloaded with lipid, increases the capacity of microglial cells to phagocytose Aβ^17^. While these studies show a clear value of ACAT1 inhibition in reducing AD pathology, the mechanisms underlying these beneficial effects have remained unclear. Specifically, the mechanism by which ACAT1 inhibition in microglia cells increases the endocytosis of Aβ to enhance degradation of Aβ has not yet been elucidated.

Triggering Receptor Expressed on Myeloid Cells 2 (TREM2) has been shown to enhance phagocytosis of cholesterol-rich substances such as myelin and cell membrane debris. In addition to phagocytosis, TREM2 signaling, mediated by DAP10/12, transcriptionally regulates genes related to cellular proliferation and survival, pro-inflammatory cytokine release, and cholesterol metabolism^17^. TREM2/DAP12 signaling is essential for the transition of microglial cells into a highly phagocytic disease-associated phenotype (DAM) which has been intensely investigated in AD^20–22^. TREM2-enriched microglia are found surrounding amyloid plaques^23^ with TREM2 capable of promoting phagocytosis ^24^. Additionally, the R47H mutation in *TREM2* is a major genetic risk factor for AD which leads to TREM2 loss of function, reducing microglial clustering around plaques^25^. Given the importance of TREM2 in microglial phagocytosis, cholesterol metabolism and AD pathogenesis, we hypothesized that impaired cholesterol storage due to inhibition of ACAT1 leads to enhanced TREM2-mediated uptake of Aβ by microglia.

Here, we find that TREM2 and low-density lipoprotein-like receptor 1 (LRP1) play central roles in the mechanism by which ACAT1 inhibition increases Aβ uptake by microglial cells. We show that enhanced uptake of Aβ by microglia following ACAT1 inhibition occurs via the cleavage of TREM2 by ADAM10/17, leading to increased release of soluble TREM2 (sTREM2) into the media. Multiple studies have reported sTREM2 binds Aβ, preventing aggregation and increasing clearance^24,26,27^. We show the addition of recombinant sTREM2 to the media of microglial cells enhances Aβ uptake to levels comparable to those observed following ACAT1 inhibition. Furthermore, when TREM2 was knocked out or TREM2 shedding was prevented with ADAM10/17 inhibition, ACAT1 inhibition no longer enhanced Aβ uptake in microglia. We also show that ACAT1 inhibition increases protein levels of LRP1. LRP1 is a widely expressed receptor, which has been previously shown in microglia to enhance Aβ uptake, clearance, and degradation^28^ while also modulating inflammatory responses ^29,30^. Here, we show that LRP1 is required for microglial sTREM2-mediated Aβ uptake, strongly suggesting LRP1 is a receptor for the internalization of the sTREM2-Aβ complex. In summary, we show that Aβ uptake is enhanced in microglia following the inhibition of ACAT1, via a mechanism in which TREM2 shedding mediated by ADAM10/17 cleavage is increased, resulting in increased uptake of Aβ in an LRP1-dependent manner.

This novel mechanism of Aβ clearance provides a unique opportunity to utilize ACAT1 inhibitors or other therapeutic methods to induce the generation of sTREM2 and, thereby, promote microglial Aβ uptake by LRP1.

## Results

### ACAT1 Inhibition Decreases Cholesteryl Esters and Increases Aβ Uptake in Mouse BV2 and Human Microglial Cells

Inhibition of ACAT1 prevents the formation of cholesteryl esters (CE) providing a measurable parameter to assess the efficacy of the ACAT1 inhibitor, Avasimibe (AV), in microglial cells^18,31^. To optimize AV treatment, we treated wild-type (WT) BV2 microglial cells with increasing concentrations of AV for 24 hours, after which CE levels were analyzed. 24-hour treatment with AV decreased CE levels at nearly all concentrations (Figure 1A) in a dose-dependent manner with 2.5 µM and 5 µM AV decreasing CE by roughly 20% and 10 µM and 15 µM leading to an approximately 30% decrease in CE levels compared to control treated cells. These findings demonstrate that AV inhibits CE production in BV2 microglia cells.

**Figure 1:**
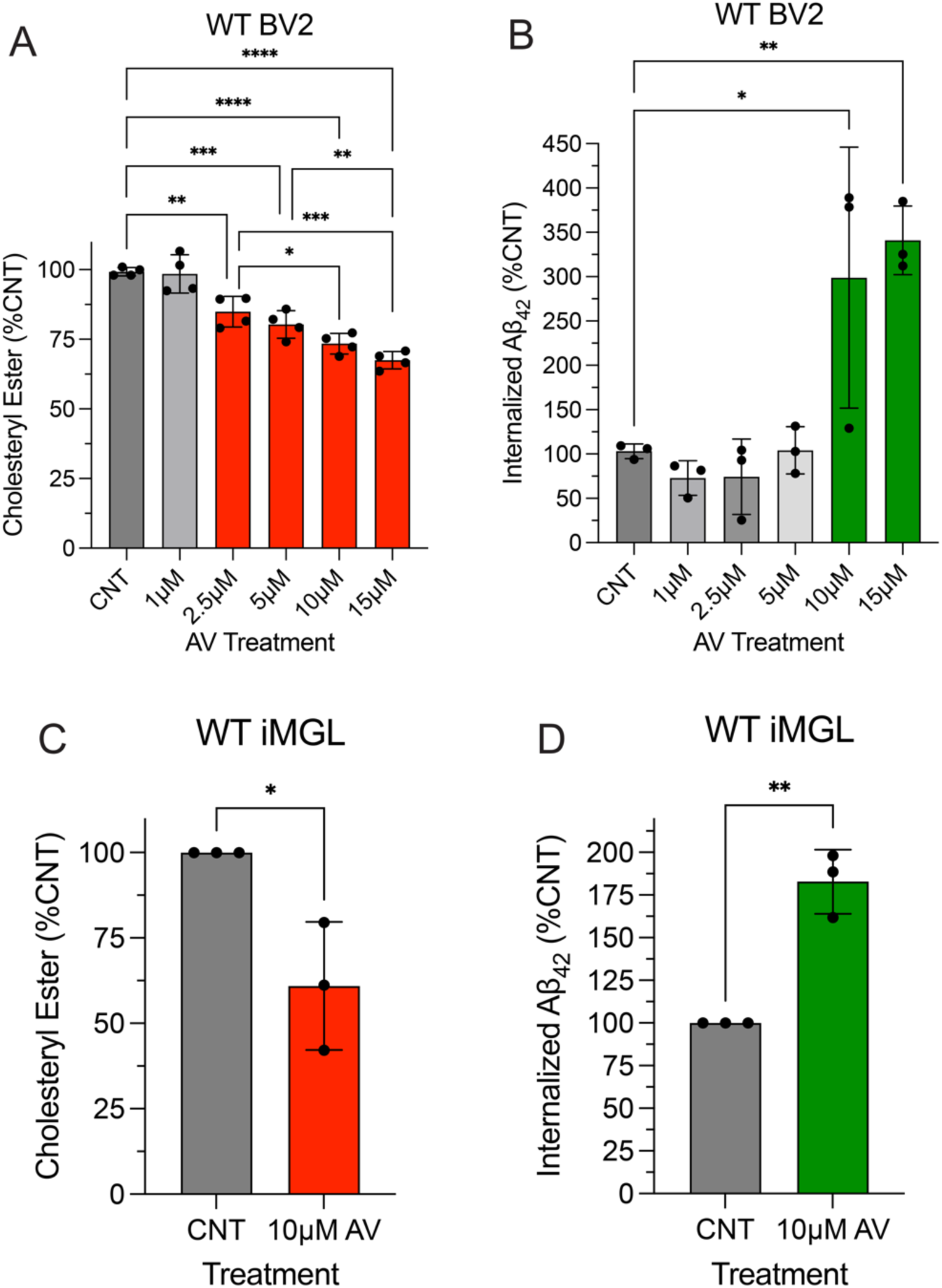
ACAT1 Inhibition Decreases Cholesteryl Ester Levels and Increases Aβ_42_ Uptake in Human and Mouse Microglial Cells. **A, B.** Mouse wild-type (WT) BV2 cells were treated with increasing concentrations of Avasimibe (AV) for 24 hours, 300 nM Aβ_42_ was added at hour 21 and incubated for the remaining three hours. **C, D** Human WT iMGL cells were treated with 10 µM AV for 24 hours, 150 nM Aβ_42_ was added at hour 21 and incubated for 1.5 hours. **A, C.** Cholesteryl ester (CE) levels were calculated by subtracting free cholesterol from total cholesterol, which was measured using a fluorometric assay and normalized to total protein. Histograms show quantification of CE normalized to total cholesterol expressed as mean ± SEM of at least three independent experiments relative to control (CNT) (* p<0.05, ** p<0.01, *** p<0.001, **** p<0.0001). **B, D** Aβ_42_ uptake was analyzed from cell lysates was quantified using an ELISA detection method, histograms show quantification as the mean ± SEM normalized to total protein of three independent experiments relative to CNT (* p<0.05 and ** p<0.01). Statistical analysis was performed using a one-way ANOVA with Tukey’s post hoc test (**A, B**) or unpaired t-test (**C, D**).

Next, we assessed the uptake of synthetic Aβ_42_ following 24 hours AV treatment in WT BV2 microglial cells. Aβ_42_ was added directly to the media of AV-treated and control cells and the cells were incubated at 37°C allowing for Aβ_42_ uptake. Internalized Aβ_42_ was then measured in cell lysates. Aβ_42_ uptake significantly increased by 195.8% and 237.9% following treatment with 10 µM and 15 µM AV, respectively (Figure 1B). In human induced pluripotent stem cell (iPSC)-derived microglial cells (iMGL), 10 µM AV treatment for 24 hours decreased CE levels by 39.1% (Figure 1C) and increased Aβ uptake by 82.8% (Figure 1D). Taken together, these data show AV treatment prevents the formation of CE by ACAT1 in both mouse BV2 and human iMGL cells, leading to increased Aβ uptake in a dose-dependent manner.

### TREM2 is Required for Increasing Aβ_42_ Uptake Following ACAT1 Inhibition in Human Microglial Cells

Next, we asked whether the increase in microglial uptake of Aβ following ACAT1 inhibition is dependent on TREM2, a myeloid-specific receptor which binds multiple ligands with high affinity, including Aβ. Binding of high-affinity ligands to TREM2 initiates multiple signaling cascades through the ITAM domain of DAP12 which leads to a diverse set of functional responses including transcriptional regulation, metabolic changes, and phagocytosis^32^. To determine whether TREM2 is required for increased Aβ uptake following ACAT1 inhibition, we first tested TREM2 knockout (TREM2 K/O) iMGLs. Figure 2A shows knockout of TREM2 significantly reduced Aβ_42_ uptake in iMGLs (by 55.0%) compared to WT iMGL cells. To determine whether TREM2 is also required for AV-mediated increased Aβ_42_ uptake in human iMGL, we treated both WT and TREM2 K/O iMGL cells with DMSO control (CNT) or 10 µM AV for 24 hours. Following AV treatment, we added 150 nM synthetic Aβ_42._ After 1.5 hours, the cells were lysed and internalized Aβ_42_ was measured. TREM2 K/O abrogated the ability of AV to augment microglial Aβ_42_ uptake as compared to WT cells (Figure 2B). WT iMGL cells treated with AV exhibited an 82.8% increase in Aβ_42_ uptake compared to control treated WT iMGLs and a 148.6% increase as compared to control treated TREM2 K/O iMGLs. However, no significant change in Aβ_42_ uptake was observed in TREM2 K/O iMGLs treated with AV compared to control treated TREM2 K/O iMGLs. When comparing WT versus TREM2 K/O iMGLs, both treated with AV, we observed a 161.0% increase in Aβ_42_ uptake in WT iMGLs versus TREM2 K/O iMGLs. Collectively, these data demonstrate that augmented uptake of Aβ_42_ by human iMGL treated with ACAT1 inhibitors is dependent on the presence of TREM2.

**Figure 2:**
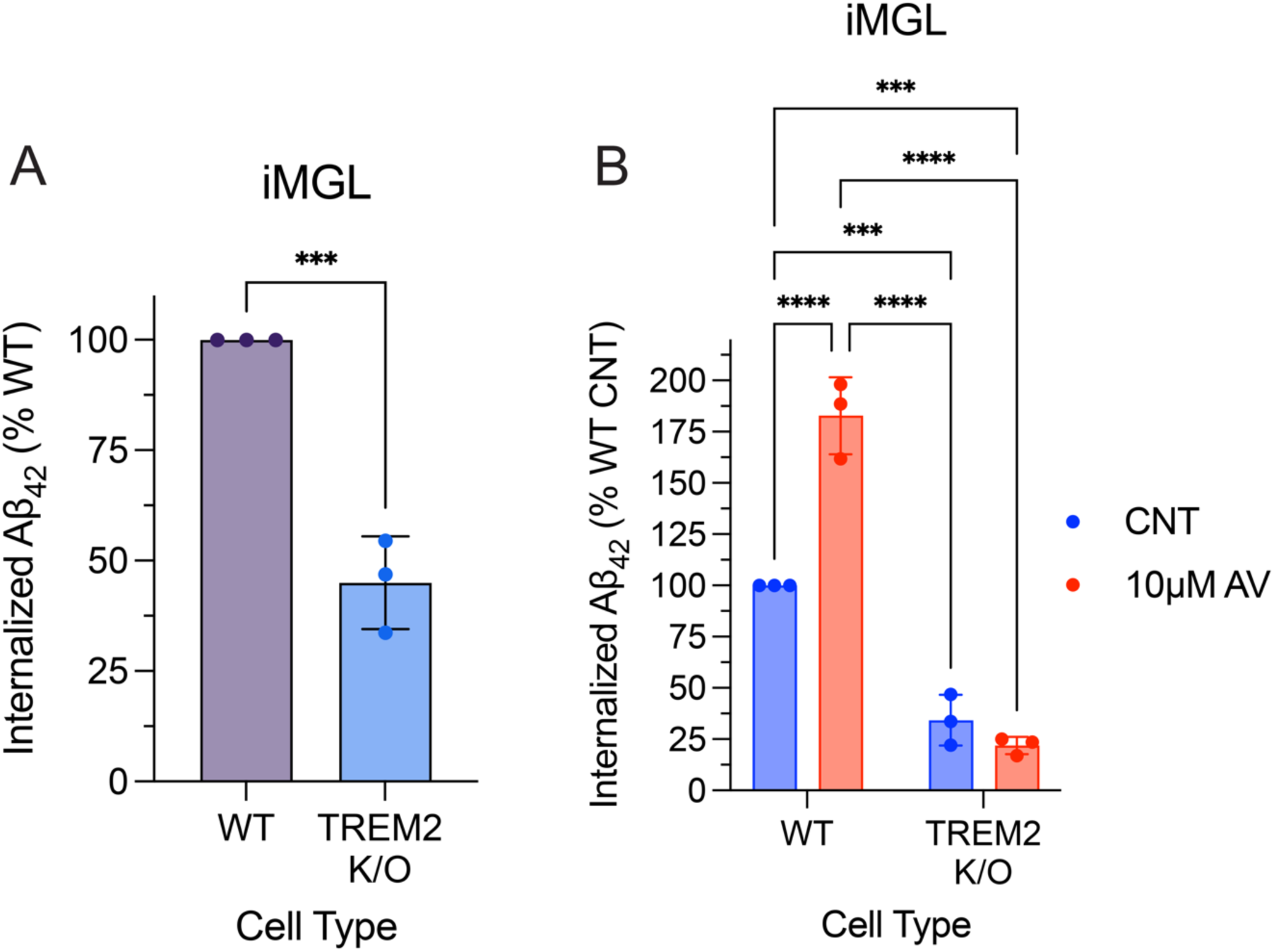
TREM2 is Required for Increased Aβ_42_ Uptake in Human iMGL following ACAT1 Inhibition. **A.** Uptake of 150 nM Aβ_42_ was measured after a 1.5-hour incubation in wild-type (WT) and TREM2 knockout (TREM2 K/O) iMGL cells. **B**. WT and TREM2 K/O iMGL cells were treated with 10 µM AV for 24 hours prior to 150 nM Aβ_42_ incubation for 1.5 hours. Aβ_42_ uptake was assessed using cell lysates, quantifying Aβ_42_ using an ELISA detection method. Histograms show quantification as the mean ± SEM (*** p<0.001, and **** p<0.0001) normalized to total protein of three independent experiments relative to WT (**A**) or WT control (CNT) (**B**). Statistical analysis was performed using a two-way ANOVA with Tukey’s post hoc test (**B**) or unpaired t-test (**A**).

### Conditioned Media of ACAT1 Inhibitor-Treated Microglia Drives Increased Aβ_42_ Phagocytosis

Next, we set out to investigate the mechanism by which TREM2 contributes to enhanced uptake of Aβ_42_ following ACAT1 inhibition. We initially hypothesized ACAT1 inhibition would lead to increased TREM2 protein levels resulting in increased phagocytosis of Aβ. However, Western blot analysis revealed that treatment of WT BV2 cells with AV, instead, resulted in decreased full-length TREM2 protein levels (44.6%) (Figure 3A and B). While initially surprising, decreasing TREM2 protein levels is a suitable physiological response to protect cells with impaired cholesterol storage from further internalizing cholesterol-rich substances.

**Figure 3:**
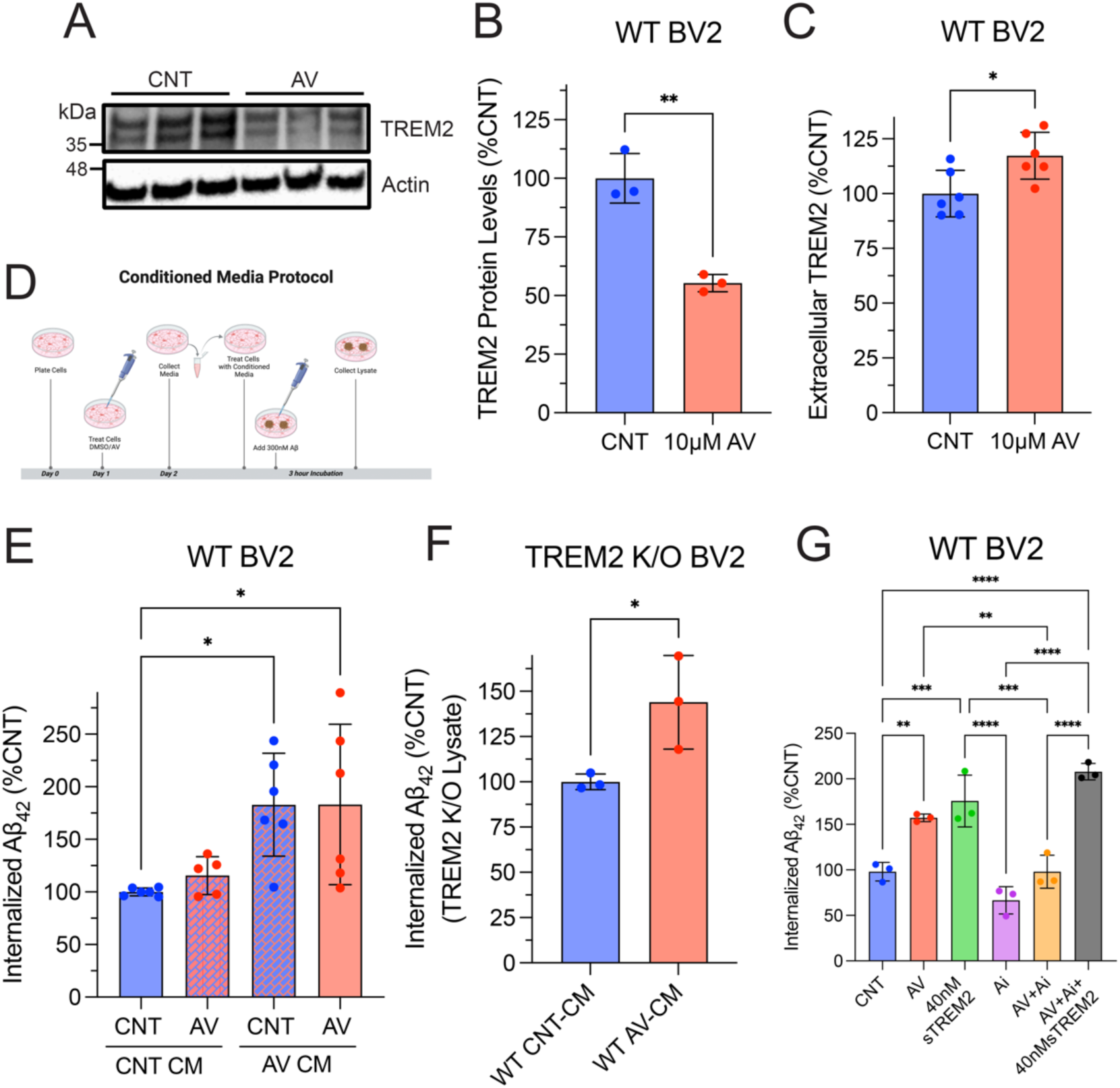
ACAT1 Inhibition Enhances Soluble TREM2 Protein Levels Which are Required for Increased Microglial Uptake of Aβ_42_. Wild-type (WT) BV2 cells treated with 10 µM AV for 24 hours. Cell lysate was collected and analyzed using SDS- PAGE. **A.** Representative western blot image of three experimental replicates measuring TREM2 and Actin in WT BV2 cells. **B** Histograms show quantification of TREM2 levels using ImageJ and normalized to Actin expressed as the mean ± SEM (** p<0.01). Statistical analysis was performed using an unpaired t-test. **C.** Extracellular TREM2 quantified using a sandwich ELISA. Histograms shows quantification of TREM2 in the media normalized to total protein from cell lysate expressed as mean ± SEM of at least three independent experiments relative to DMSO control treated cells (CNT) (*p<0.05). Statistical analysis was performed using an unpaired t-test **D.** Schematic of Conditioned Media (CM) experimental design. Media is collected from WT BV2 cells treated with 10 µM Avasimibe (AV) or DMSO (CNT) after 24 hours then place on new cells and treated with 300 nM Aβ_42_ and incubated for three hours. **E**. WT BV2 cells treated with CNT or AV for 24 hours were then treated with CNT or AV CM immediately before a three-hour 300 nM Aβ_42_ incubation. **F.** TREM2 K/O BV2 cells were treated with WT CNT-CM or WT AV-CM immediately before a three-hour 300 nM Aβ_42_ incubation **G**. WT BV2 cells treated with DMSO (CNT), 10 µM AV, or 3 µM ADAM10/17 inhibitor (GW280264X) (Ai) for 24 hours followed by a three-hour incubation with 300 nM Aβ_42_ and 40 nM sTREM2 as stated. **E-G**. Aβ_42_ uptake was measured from cell lysate using an ELISA based assay. Histograms show quantification of internalized Aβ_42_ normalized to total protein expressed as mean ± SEM of at least three independent experiments relative to CNT/CNT CM (**E**), CNT (**F**), WT CNT-CM (**G**) (* p<0.05, ** p<0.01, *** p<0.001, and **** p<0.0001). Statistical analysis was performed using a one-way ANOVA with Tukey’s post hoc test (**E,G**) or unpaired t- test (**F**)

We next explored how a decrease in full-length TREM2 affects Aβ uptake in AV-treated microglial cells with impaired cholesterol storage. Multiple prior studies have shown the extracellular soluble TREM2 (sTREM2) fragment, produced by cleavage and shedding of full-length membrane TREM2, binds Aβ, prevents fibrilization, and enhances Aβ phagocytosis^33^. Using a sandwich ELISA, we measured extracellular levels of sTREM2 in the cell culture medium following AV treatment. sTREM2 levels were significantly increased (17.3%) compared to control treated cells (Figure 3C). Thus, ACAT1 inhibition promotes cleavage of membrane TREM2 and shedding of sTREM2 into the culture medium, leading to the hypothesis that it then binds Aβ and enhances microglial uptake.

To test this hypothesis, we first asked whether conditioned media (CM) from AV-treated WT BV2 microglia cells is necessary and sufficient to increase microglial Aβ_42_ uptake. Figure 3D shows a schematic of the experimental design for the CM experiments (also described in the methods section). Briefly, WT BV2 cells were treated with 10 µM AV or DMSO (CNT) for 24 hours. CM from these cells was then collected and added to new cells which had been treated with either 10 µM AV or DMSO (CNT) for 24 hours. Immediately following the replacement of media, 300 nM Aβ_42_ was added to the cells and incubated for three hours. This resulted in four treatment conditions, for which we measured Aβ_42_ uptake in WT BV2 cells: 1. CNT treated cells incubated with CNT-CM (CNT CNT-CM), 2. AV-treated cells incubated with CNT-CM (AV CNT-CM), 3. CNT treated cells incubated with AV CM (CNT AV-CM), and 4. AV treated cells incubated with AV-CM (AV AV-CM).

Analysis of Aβ_42_ uptake revealed AV-CM significantly increased Aβ_42_ uptake in CNT treated cells (82.9%) and AV treated cells (83.1%) compared to WT BV2 cells which were not treated with AV (CNT CNT-CM) (Figure 3E). These findings indicate that CM from AV-treated cells is sufficient for augmenting Aβ_42_ uptake in microglia. Meanwhile, WT BV2 cells treated with CNT or AV overnight, for which media was then replaced with CNT-CM, showed no significant increase in Aβ_42_ uptake. (Figure 3E). These data indicate AV-CM is necessary for inducing increased uptake of Aβ_42_ in microglia. Collectively, these findings demonstrate that factors released into the CM by AV-treated microglia are both sufficient and necessary to increase microglial Aβ_42_ uptake following ACAT1 inhibition. Furthermore, these data suggest that depletion of intracellular CE levels following ACAT1 inhibition with AV is responsible for the elevation in soluble factors that are released by microglia and enhance Aβ_42_ uptake.

### Increased Levels of sTREM2 in the Conditioned Media of ACAT1 Inhibitor-Treated Microglia Drive Enhanced Microglial Aβ_42_ Phagocytosis

We next investigated the role of increased levels of sTREM2 in the conditioned media of AV-treated microglia in augmented microglial Aβ_42_ uptake. For this purpose, we first analyzed Aβ_42_ uptake in TREM2 K/O BV2 cells treated with WT CNT-CM or WT AV-CM. WT AV-CM significantly increased (43.9%) Aβ_42_ uptake in TREM2 K/O BV2 cells compared to TREM2 K/O cells treated with WT CNT-CM (Figure 3F). These data prompted us to further test whether sTREM2 is a necessary component in AV-CM that drives enhanced Aβ_42_ uptake in microglia. For this purpose, we investigated whether preventing the cleavage of TREM2 by its major sheddases, ADAM10/17, using the ADAM10/17 inhibitor, GW280264X, abrogates ACAT1 inhibitor-mediated augmentation of microglial Aβ_42_ uptake. Figure 3G shows inhibition of ADAM10/17 with GW280264X, alone, does not significantly change Aβ_42_ uptake compared to control treated cells. However, when GW280264X treatment was combined with AV treatment, Aβ_42_ uptake was no longer increased compared to control treated cells; AV treatment alone increased uptake by 57.1% (Figure 3G). These results indicate that ADAM10/17 activity, alone, does not significantly contribute to baseline Aβ_42_ uptake but is necessary for enhancement of microglial Aβ_42_ uptake following inhibition of ACAT1.

Having shown cleavage of TREM2 by ADAM10/17 is necessary for the enhancement of microglial Aβ_42_ uptake following inhibition of ACAT1, we next asked whether supplementation of the media with recombinant sTREM2 protein could rescue the effects of ADAM10/17 inhibition using GW280264X, on AV-enhanced microglial Aβ_42_ uptake. The addition of 40 nM sTREM2 to WT BV2 cells significantly increased Aβ_42_ uptake by 75.6% compared to control treated cells; this is comparable to the 57.2% increase in Aβ_42_ uptake that was observed with AV treatment (Figure 3G). Importantly, 40 nM sTREM2 also rescued the effect of ADAM10/17 inhibition in AV-treated WT BV2 cells, significantly increasing Aβ_42_ uptake by 107.9% compared to control treated cells (Figure 3G). Taken together, these data strongly suggest that the release of sTREM2 via ADAM10/17-mediated cleavage of membrane TREM2, is the major driver of increased Aβ uptake following ACAT1 inhibition in microglia.

### LRP1 is Required for Enhanced sTREM2-Mediated Aβ_42_ Uptake following ACAT1 Inhibition in Microglia

sTREM2 has previously been shown to bind Aβ and prevent oligomerization and fibrillation as well as increase Aβ phagocytosis^33^. However, the mechanism by which sTREM2 increases internalization of Aβ has remained unknown. LRP1 is a well-known receptor important for lipid metabolism^34^, protein clearance^35^, and cell signaling^36,37^ and has been previously implicated in AD pathology^38,39^, based on binding Aβ, and promoting rapid endocytosis and clearance^28^. Additionally, LRP1 regulates intracellular signaling events involved in lipid metabolism, cholesterol storage, and fatty acid synthesis^34^. Thus, we next investigated whether LRP1 plays a role in sTREM2-dependent augmentation of microglial Aβ uptake following ACAT1 inhibition.

First, we observed WT BV2 cells treated with 10 µM AV for 24 hours exhibit significantly increased protein levels of LRP1 (102.7%) compared to DMSO CNT treated cells (Figure 4A and B). In Figure 4C we show Aβ_42_ uptake in LRP1 K/O BV2 cells was significantly decreased (61.92%) as compared to WT BV2. Interestingly, ACAT1 inhibition was unable to increase Aβ_42_ uptake in LRP1 K/O BV2 microglial cells as compared to control cells (Figure 4C). Taken together, these data indicate that LRP1 is required for AV-mediated enhancement of microglial Aβ_42_ uptake.

**Figure 4:**
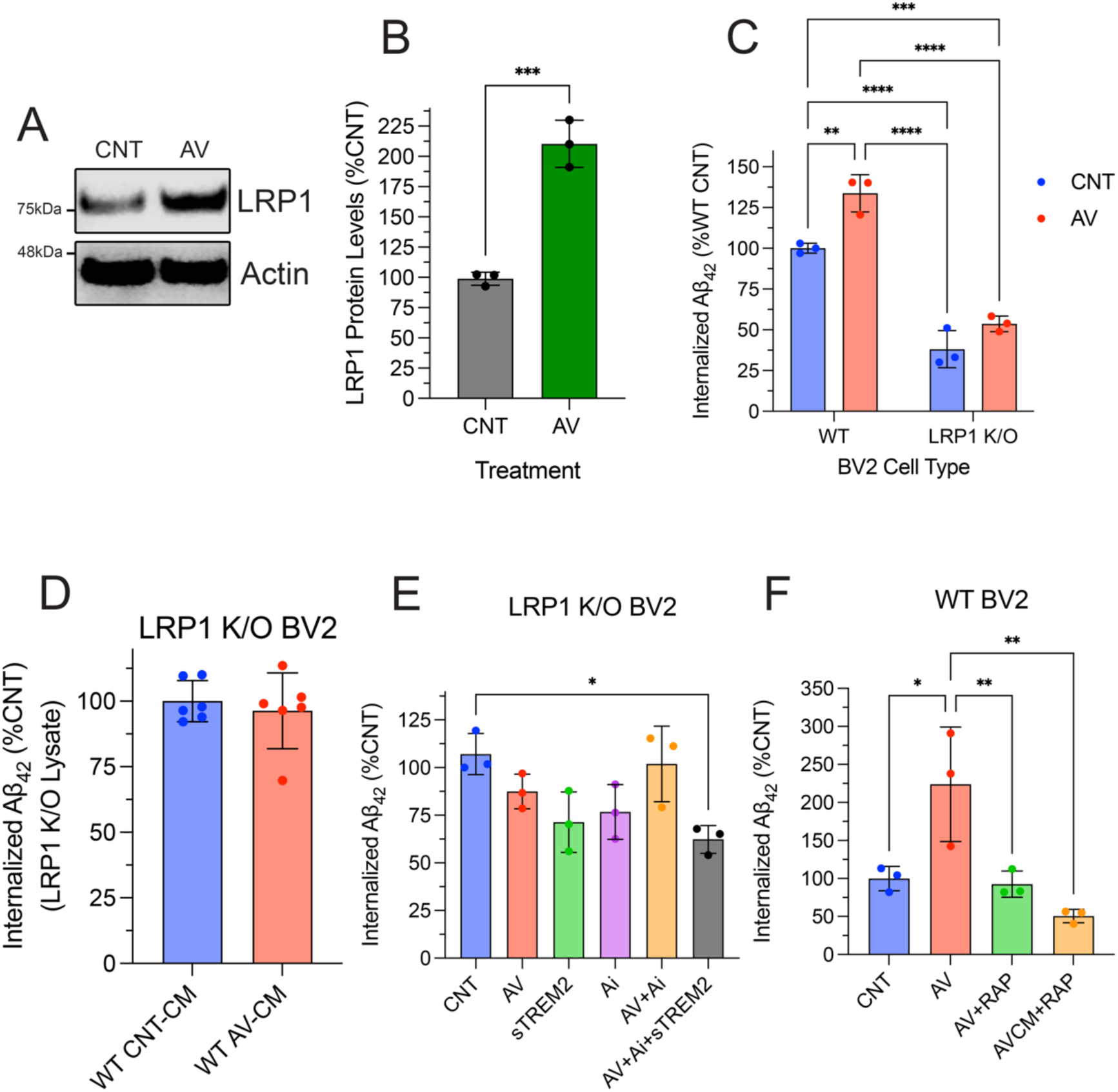
LRP1 is a Receptor for the sTREM2:Aβ Complex and is Required for Increased Microglial Aβ Uptake Following ACAT1 Inhibition. Wild-type (WT) BV2 cells treated with 10 µM Avasimibe (AV) for 24 hours. Cell lysate was collected and analyzed using SDS-PAGE. **A.** Representative western blot image measuring LRP1 and Actin in WT BV2 cells. **B**. Histogram shows quantification of at least three independent experiments expressed as the mean ± SEM (*** p<0.001). Statistical analysis was performed as an unpaired t-test. **C**. WT and LRP1 knockout (LRP1 K/O) BV2 cells were treated with DMSO (CNT) or 10 µM AV for 24 hours followed by a three- hour incubation with 300 nM Aβ_42._ Quantification of internalized Aβ_42_ was performed on cell lysate using an ELISA based assay normalized to total protein levels expressed as the mean ± SEM of at least three independent experiments relative to WT CNT (** p<0.01, *** p<0.001, and **** p<0.0001). Statistical analysis was performed using a two-way ANOVA with Tukey’s post hoc test. **D**. Quantification of internalized Aβ_42_ from LRP1 K/O cells treated with CNT or AV conditioned media (CM) from WT BV2 cells of at least three independent experiments as the mean ± SEM. Statistical analysis was performed as an unpaired t-test. **E-F**. WT BV2 cells treated with stated conditions were compared to CNT Aβ_42_ uptake. Histograms show quantification of internalized Aβ_42_ normalized to total protein expressed as the mean ± SEM of at least three independent experiments relative to CNT (* p<0.05, ** p<0.01, *** p<0.001, and **** p<0.0001). Statistical analysis was performed using an or one-way ANOVA with Tukey’s post hoc test.

Knockout of LRP1 could also impact protein levels of other receptors involved in cholesterol metabolism, e.g. TREM2, which would limit the amount of sTREM2 released and AV-induced Aβ_42_ uptake. To address this possibility, we measured TREM2 protein levels in LRP1 K/O BV2 cells and observed significantly lower levels of total TREM2 levels as compared to WT BV2 cells. However, when LRP1 K/O BV2 cells were treated with AV, sTREM2 levels were increased by 16.3% as compared to DMSO (CNT) (Supplemental Figure 2). This increase was similar to that observed in WT BV2 cells (17.3% in Figure 3C). We also found that AV-CM from WT BV2 cells did not increase Aβ_42_ uptake in LRP1 K/O BV2 cells (Figure 4D). Collectively, these data suggest that LRP1 is a crucial receptor for augmented microglial uptake of Aβ_42_ following ACAT1 inhibition.

Next, we treated LRP1 K/O BV2 cells with the same combinations of GW280264X, AV, and recombinant sTREM2 as the WT BV2 cells shown in Figure 4E. LRP1 K/O BV2 cells continued to exhibit no change in Aβ_42_ uptake when treated with 10 µM AV for 24 hours. Additionally, we observed no change in Aβ_42_ uptake in LRP1 K/O BV2 cells following treatment with 40 nM recombinant sTREM2. Next, as observed in the WT BV2 cells, LRP1 K/O BV2 cells treated with ADAM10/17 inhibitor, GW280264X, alone or combined with AV treatment, revealed no change in Aβ_42_ uptake. Finally, treatment with recombinant 40 nM sTREM2 led to no change in Aβ_42_ uptake in LRP1 K/O cells treated with both GW280264X and AV, in contrast to the results from WT BV2 cells. Taken together, these data suggest that following ACAT1 inhibition, LRP1 most likely serves as a major receptor for microglial uptake of Aβ_42_, via internalization of an sTREM2-Aβ complex.

To further validate the requirement of LRP1 in AV-enhanced microglial Aβ uptake, we tested the effects of Receptor Associated Protein (RAP), a well-known LRP1 antagonist that binds LRP1 and reduces its ligand binding capacity ^35,40^. In WT BV2 cells treated with 250 nM RAP 30 minutes prior to the addition of Aβ_42,_ we observed a significant decrease in Aβ_42_ uptake (Supplemental Figure 1B). Next, we treated WT BV2 cells with RAP following 24-hour AV treatment and 30 minutes prior to the addition of Aβ_42_. Figure 4F shows AV-mediated enhancement of Aβ_42_ uptake was abrogated in the presence of RAP as compared to control treated cells. Additionally, when control WT BV2 cells were treated with WT BV2 AV-CM treatment together with RAP, AV-enhanced Aβ_42_ uptake was abrogated compared to control WT BV2 cells (Figure 4F). Collectively, these findings show that LRP1 is necessary for AV-induced enhancement of Aβ uptake in microglia and that the observed effects of decreased Aβ uptake in LRP1 K/O BV2 cells is not due to decreased TREM2 levels.

Overall, our combined findings support a mechanism for ACAT1 inhibitor-enhanced microglial uptake of Aβ, wherein altered cholesterol metabolism due to ACAT1 inhibition leads to elevated shedding of sTREM2, via increased ADAM10/ADAM17-mediated cleavage of membrane TREM2. ACAT1 inhibition simultaneously increases levels of LRP1, which serves to phagocytose Aβ bound to sTREM2 (Figure 5).

**Figure 5:**
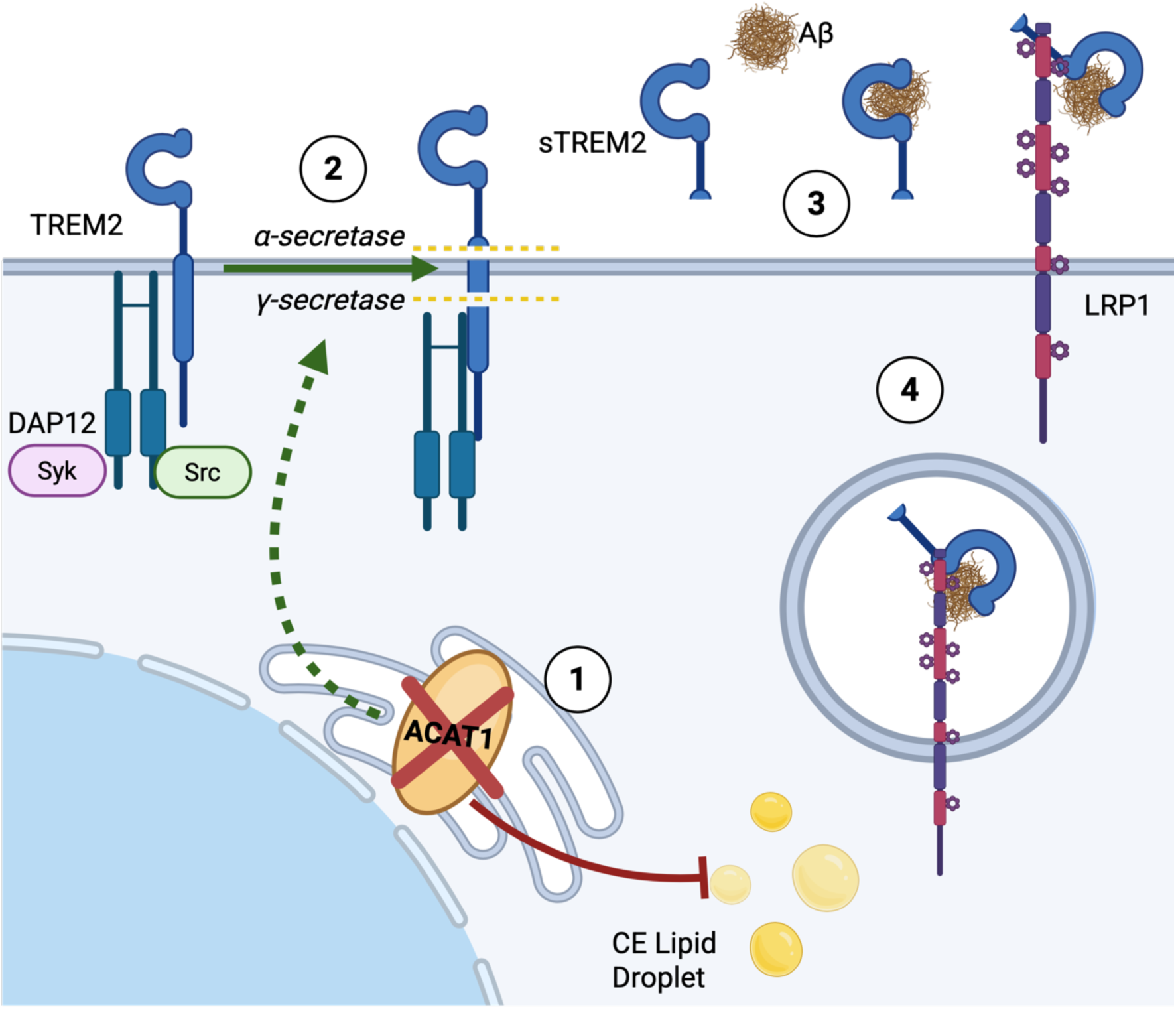
Model for How ACAT1 Inhibition Increases Aβ Uptake in Microglia via Increased Shedding of sTREM2 by α-Secretase and Expression of LRP1. Diagram of the proposed mechanism showing the role of TREM2 in ACAT1 inhibition enhanced Aβ uptake in microglia. 1. ACAT1 inhibition prevents cholesteryl ester (CE) formation leading to decreased CE levels and an accumulation of free cholesterol. 2. TREM2 shedding is increased through α-secretase (ADAM10/17). 3 Increased levels of sTREM2 in the media bind Aβ. 4. The sTREM2-Aβ complex binds LRP1 whereupon the complex is internalized resulting in increased Aβ uptake.

## Discussion

ACAT1 inhibition has been previously shown to reduce multiple aspects of AD pathology *in vivo*^16^ and *in vitro*^18^. Here we show ACAT1 inhibition increases Aβ uptake in both human iMGL and mouse BV2 microglial cells. Furthermore, we show that the absence of TREM2 in iMGL cells significantly impairs augmented Aβ uptake following ACAT1 inhibition. TREM2 plays an essential role in maintaining microglial metabolic fitness during stress events^41^ and is necessary for microglial cells to fully transition into the disease-associated microglia (DAM) phenotype which is sustained in the presence of Aβ- induced pathology. Furthermore, TREM2 regulates cholesterol transport and metabolism through transcriptional regulation and endocytosis of cholesterol rich substances such as myelin and cell debris^42^. For these reasons, we and others have proposed that TREM2 plays a major role in sensing and employing cellular changes necessary to adapt to impaired cholesterol storage, supporting a connection between TREM2 activity and ACAT1 inhibition^42,43^.

We provide further support for a role of TREM2 in enhanced microglial Aβ uptake following ACAT1 inhibition by showing decreased full-length TREM2 protein levels in AV-treated microglial cells undergoing impaired cholesterol storage (Figure 3). Previous studies have shown that cholesterol depletion enhances the activity of ADAM10/17, leading to increased shedding of substrates such as the interleukin-6 receptor (IL-6)^44^. Here, we observed that ACAT1 inhibition enhanced ADAM10/17- mediated cleavage of TREM2 leading to increased shedding of sTREM2 into the media of AV-treated microglial cells (Figure 3A). This finding led us to propose a novel mechanism for enhanced Aβ uptake following ACAT1 inhibition, wherein sTREM2 serves as a key mediator in this process. In support of this hypothesis, we showed that media conditioned by WT BV2 microglial cells treated with an ACAT1 inhibitor was sufficient to increase Aβ uptake in WT BV2 cells, which had not been treated with an ACAT1 inhibitor (Figure 3E). Importantly, when we replaced the media of ACAT1 inhibitor-treated WT BV2 microglial cells with conditioned media from control treated WT BV2 cells, Aβ uptake was no longer increased (Figure 3E). In further support of our hypothesis that sTREM2 is the major secreted factor enhancing Aβ uptake, we show that ADAM10/17 inhibition abrogated ACAT1 inhibitor-mediated enhancement of Aβ uptake (Figure 3G). Moreover, the effect of ADAM10/17 inhibition on ACAT1 inhibitor-mediated Aβ uptake could be rescued by the simple addition of recombinant sTREM2 during Aβ incubation. Collectively, these data establish that the shedding of sTREM2 is the major driver of enhanced microglial uptake of Aβ following inhibition of ACAT1.

While the role of sTREM2 in AD is complex and not fully understood^45,46^, sTREM2 has been previously shown to prevent Aβ aggregation and promotes phagocytosis^27,46^. However, the mechanism by which sTREM2 promotes Aβ uptake has remained unclear. Here, we show LRP1 is essential for enhanced Aβ uptake mediated by sTREM2 following ACAT1 inhibition. We show this through multiple experiments demonstrating 1. ACAT1 inhibition does not increase Aβ uptake in LRP1 K/O BV2 cells (Figure 4C), 2. Conditioned media from WT BV2 cells treated with ACAT1 inhibitor does not increase Aβ uptake in LRP1 K/O cells (Figure 4D), 3. The addition of sTREM2 was unable to increase Aβ in LRP1 K/O cells (Figure 4E) and 4. WT BV2 cells treated with the potent LRP1 antagonist, RAP, no longer display enhanced Aβ uptake following ACAT1 inhibitor treatment or with conditioned media from cells treated with an ACAT1 inhibitor (Figure 4F). Collectively these data strongly support LRP1 as a necessary and sufficient receptor for sTREM2-dependent, augmentation of Aβ uptake following ACAT1 inhibition in microglial cells. LRP1 is a ubiquitously expressed protein which binds many ligands and is known to function in a context-dependent manner, displaying distinct functions in different tissues and cell types^47,48^. To our knowledge, the data presented are the first to reveal LRP1 as the receptor for the sTREM2-Aβ complex.

This body of evidence robustly supports a novel and heretofore unknown mechanism for the manner in which ACAT1 inhibition leads to increased microglial phagocytosis of Aβ (Figure 5) as follows: 1. AV treatment inhibits ACAT1, thereby decreasing cholesteryl ester levels, 2. Depletion of cholesterol enhances ADAM10/17 cleavage of membrane TREM2 to promote TREM2 shedding, and 3. Elevated levels of sTREM2 in the media promotes Aβ uptake via LRP1-mediated endocytosis. This novel mechanism suggests possibility of employing ACAT1 inhibitors as TREM2 modulating therapies for enhancing microglial clearance of Aβ in a potentially safer and more affordable manner than approved anti-amyloid immunotherapies. In addition to plaque and tangle pathology, neuroinflammation and vascular pathology^49^ also play significant roles in the pathogenesis of AD. ACAT1 inhibitors have previously been shown to decrease atherosclerotic plaques^50,51^ to improve vascular function and to impact inflammatory responses^52^ via TLR4^13^ and NLRP3. These findings, together with our current data and the known ability of ACAT1 inhibitors to reduce Aβ plaque formation^16^ and promote degradation of tau^19,53^ and Aβ^18^ would suggest that ACAT1 inhibitors carry great potential as multi-modal disease modifying therapeutics for the treatment and prevention of AD.

## Acknowledgments

We thank Dr. Doo Yeon Kim and Dr. Joseph Park for providing the iMGL TREM2 Knockout cell line. We thank Donna Romano, Ana Griciuc, Dominika Pilat, Raja Bhattacharyya, Luisa Quinti, Junseok Bae, and other current and former members of the Genetics and Aging Research Unit at Massachusetts General Hospital. This research was supported by grants from the JPB Foundation (R.E.T.), Cure Alzheimer’s Fund (R.E.T.), and NIH (T32 AG000222-31 to M.J.H.). Research reported in this publication was supported by the National Institute On Aging of the National Institutes of Health under Award Number T32AG000222. The content is solely the responsibility of the authors and does not necessarily represent the official views of the National Institutes of Health

## Author Contributions

M.J.H Conceptualized project and methodology, investigation, planned and performed experiments, analyzed data, and wrote the manuscript

R.E.T. Conceptualized project and methodology, planned and supervised the study, wrote the manuscript, and acquired funding

A.M.H methodology and edited the manuscript

---

**Supplemental Figure 1:**
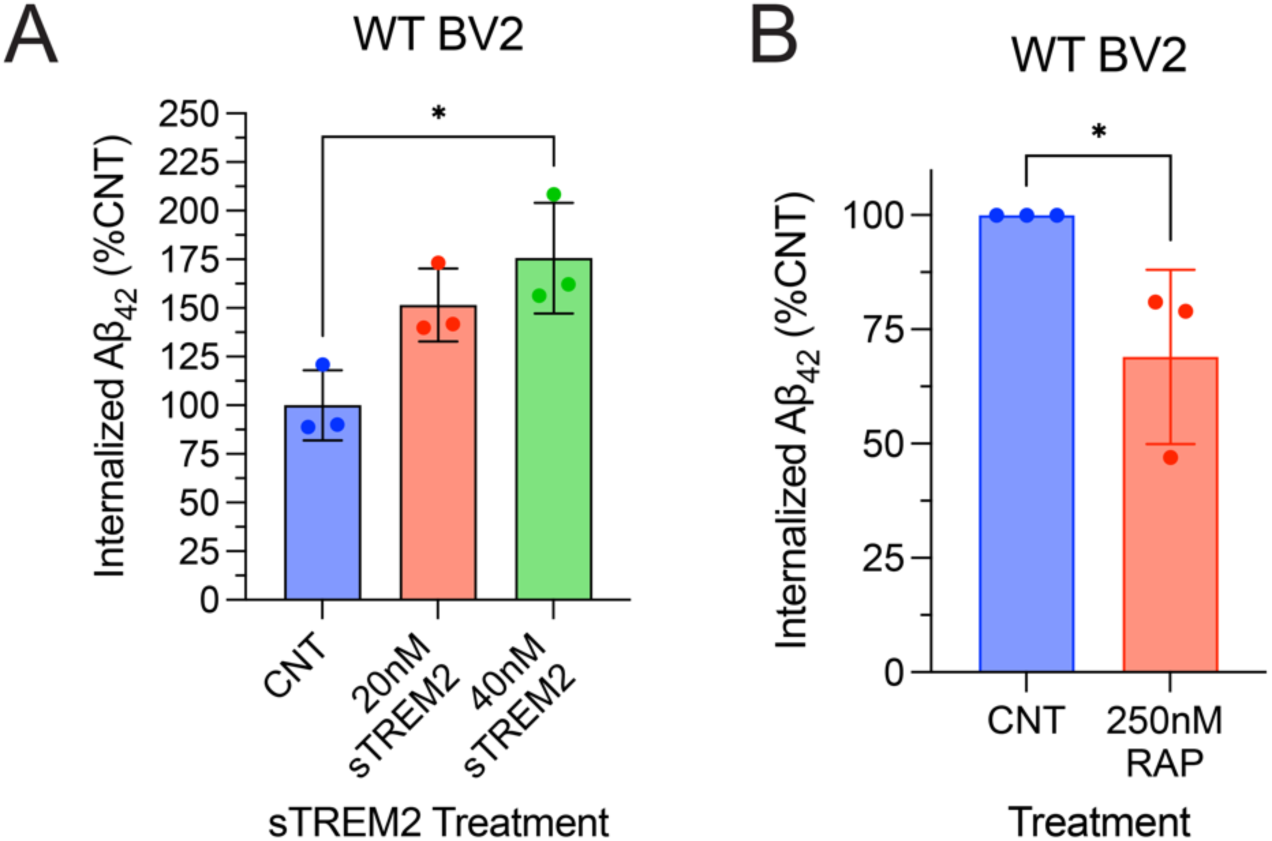
Treatment with sTREM2 Increases Aβ Uptake while RAP Decrease Aβ Uptake in WT BV2 Cells. Wild-type (WT) BV2 cells were treated with mouse recombinant sTREM2 (**A**) or 250 nM RAP (**B**) for three hours along with 300 nM Aβ_42_. Aβ_42_ uptake was measured from cell lysate using an ELISA based assay. Histogram shows quantification of internalized Aβ_42_ normalized to total protein expressed as the mean ± SEM of at least three independent experiments relative to an untreated control (CNT) (*p<0.05). Statistical analysis was performed using a one-way ANOVA (**A**) with Tukey’s post hoc test or unpaired t-test (**B).**

**Supplemental Figure 2.**
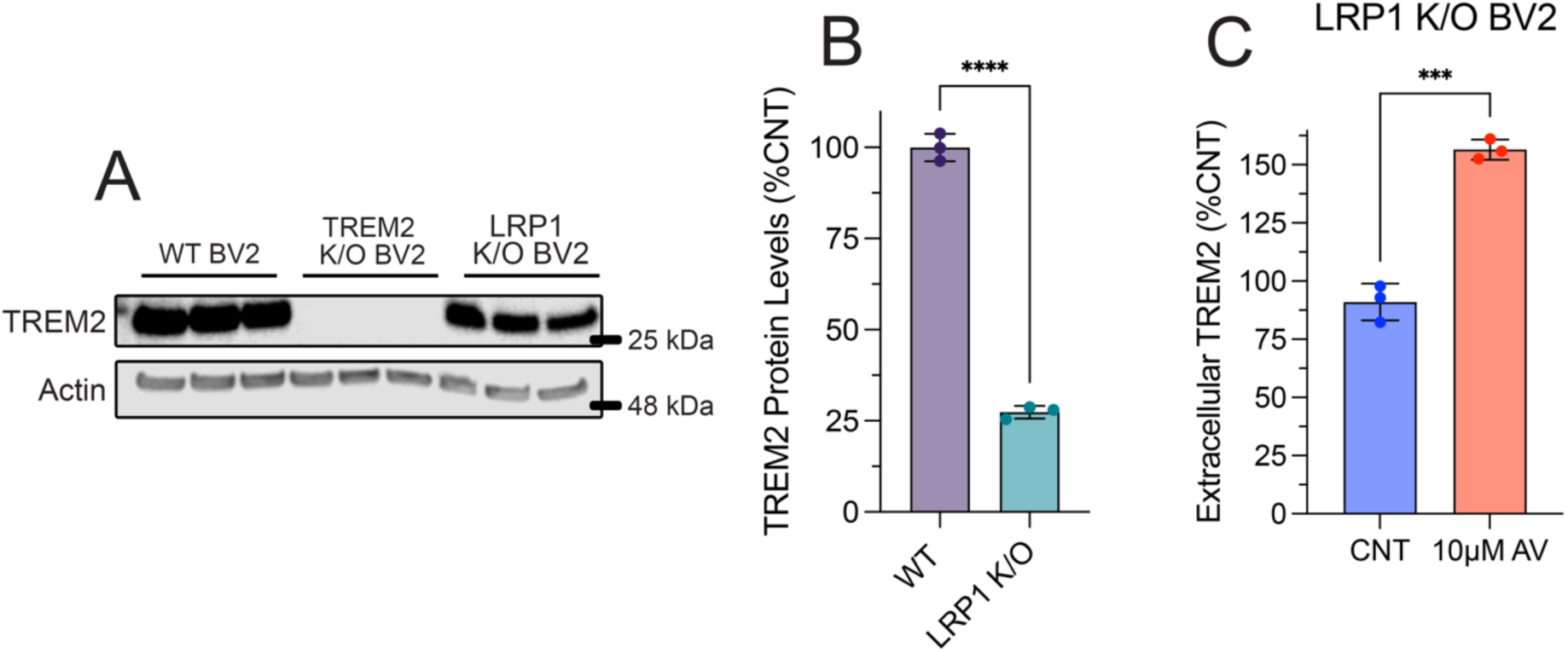
Extracellular TREM2 is Increased in LRP1 K/O BV2 Cells Treated with AV Despite Lower Total TREM2 Levels. **A**. Representative western blot image measuring TREM and Actin in wild-type (WT), TREM2 knockout (TREM2 K/O), and LRP1 knockout (LRP1 K/O) BV2 cells. **B**. Histogram shows quantification of at least three independent experiments as the mean ± SEM (**** p<0.0001). Statistical analysis was performed as an unpaired t-test. **C.** Extracellular TREM2 quantified in media from WT and LRP1 K/O cells treated with DMSO (CNT) or 10 µM Avasimibe (AV) for 24 hours using a sandwich ELISA. Histogram shows quantification of TREM2 in the media normalized to total protein from cell lysate expressed as mean ± SEM of at least three independent experiments relative to WT CNT (*p<0.05, *** p<0.001, and **** p<0.0001). Statistical analysis was performed using a one-way ANOVA with Tukey’s post hoc test

## Methods

### Mouse BV2 Microglial Cell Culture

BV2 microglia cells were maintained in Dulbecco’s modified Eagle Medium (DMEM) supplemented with 10% fetal bovine serum, 2 mM L-glutamine, and 1% antibiotics (10,000 U/mL Penicillin-Streptomycin, Gibco™) at 37°C in a humidified 5% CO_2_ incubator. Cells were passaged at 80% confluency using the dissociation agent StemPro™ Accutase™.

### Human iPSC cell culture

The parental human iPSC line was purchased from the NINDS Human Cell and Data Repository (Cell line ID - ND41865, RUID - NN0003920, male, 64 years old). Cells were routinely maintained in mTeSR™ Plus medium (100-0276, STEMCELL Technologies Inc) supplemented with 100 U/mL penicillin and 100 μg/mL streptomycin (15140122, Life Technologies) on Matrigel-coated (1% v/v) tissue culture dishes under standard conditions at 37°C in a humidified 5% CO_2_ incubator. The medium was replaced daily, and cultures were monitored for spontaneous differentiation. Standard passaging of iPSC lines was performed at 70-80% confluency using enzyme-free dissociation reagent ReLeSR™ (STEMCELL Technologies Inc) according to the manufacturer’s protocol. To achieve single cell suspensions, iPSCs were dissociated with StemPro™ Accutase™ (A11105-01, Life Technologies) for 15-20 minutes at 37°C.

### Human iMGL differentiation

Human iPSCs were differentiated to iMGL according to the protocol previously published by McQuade 2018, with minor adjustments^54,55^. The initial differentiation from iPSCs to hematopoietic progenitor cells (HPCs) was achieved using Stem Cell Technologies STEMdiff™ Hematopoietic Kit (Catalog # 05310) following the manufacturer’s protocol. In brief, on day -1, 70% confluent human iPSCs are passaged with ReLeSR (STEMCELL Technologies Inc) into mTeSR™ Plus medium onto Matrigel coated (1% v/v) 6-well plates. Small aggregates of ∼100 cells each are plated at 1-10 aggregates/cm^2^. On day 0 medium was replaced with medium A (Basal medium plus Supplement A). On day 2 50% of the medium A was replaced with fresh medium A. On day 3, all media was removed and replaced with medium B (Basal medium plus supplement B). On days 5, 7, and 10 half the medium was replaced with fresh medium B. On day 12, non-adherent cells were collected by centrifugation at 300 × g for 5 minutes. On day 0 of the iMGL differentiation ∼30K/cm^2^ HPCs were plated on (1% v/v) Matrigel-coated 100 mm dishes in 6 mL iMGL basal medium (DMEM-F12, 2X B27, 0.5X N2, 1X Glutamax, 1X NEAA, 400 µM monothioglycerol, 5 µg/mL insulin, 2X insulin-transferrin-selenium, 100 U/mL penicillin and 100 μg/mL streptomycin, and 0.05% BSA) supplemented with 100 ng/mL IL-34, 100 nM IDE1, and 25 ng/mL M-CSF. On days 2, 4, 6, 8, and 10, 3 mL basal medium plus freshly thawed tri-cytokine cocktail was added to the cells. On day 12, media was collected, leaving 3 mL of conditioned medium in the dish, and non-adherent cells were centrifuged at 300 × g for 5 minutes. The pelleted cells were returned to the dish and supplemented with fresh basal medium with tri-cytokine cocktail. On days 14, 16, 18, 20, 22, and 24 3 mL basal medium plus tri-cytokine cocktail was added to the cells. On day 25, media was collected, leaving 3 mL of conditioned medium in the dish, and non-adherent cells were centrifuged at 300 × g for 5 minutes.

Cells were resuspended in basal media supplemented with 100 ng/mL IL-34, 100 nM IDE1, 25 ng/mL M-CSF, 100 ng/mL CD200 and 100 ng/mL CX3CL1 to further mature microglia. On day 27, iMGLs were supplemented with basal medium and the five-cytokine cocktail, after which they were matured and ready for functional analyses.

### TREM2 Knockout Generation

The control hiPSC line (NDS00159) was electroporated to generate TREM2 knockout clones. Briefly, a chemically modified sgRNA (5’-CACAACACCACAGUGUUCCA-3’) targeting exon 2 of TREM2 (Transcript ID: ENST00000373113.8, chromosomal location chr6:41,161,583) was complexed with Cas9 protein to form ribonucleoprotein (RNP) complexes. The RNP complexes were delivered to iPSCs via optimized electroporation parameters. As a positive control, cells were also transfected with RELA-targeting sgRNA. After transfection, cells were recovered for 2 days in mTeSR™ Plus medium before analysis. All experiments were performed at Synthego.

Editing efficiency was assessed by PCR amplification of the targeted region using forward (5’- GGTAGAGACCCGCATCATGG-3’) and reverse (5’-CCATCCGCTCCCAACTTGTA-3’) primers, followed by Sanger sequencing. Sequence chromatograms were analyzed using Synthego’s Inference of CRISPR Edits (ICE) software to determine indel frequency.

Single cells from the edited pool were isolated by single-cell dilution for clonal expansion. Wells were imaged every 2-3 days to ensure true clonality. No selection agents were used during the clonal isolation process. Expanded clones were genotyped by Sanger sequencing, and two homozygous knockout clones (B6 and E3) were selected for further characterization. Both selected clones contained a +1 insertion at the Cas9 cleavage site, creating frameshift mutations that disrupted the TREM2 coding sequence.

The selected clones were expanded to passage 12 for final characterization and quality control. Wild-type control cells were generated through mock transfection and processed in parallel through passage 6. All cell lines tested negative for mycoplasma contamination. The final cell lines were cryopreserved at a concentration of 0.5×10⁶ cells per vial. A QC report from Synthego is provided.

### Cell Treatments

Cells were sub-cultured and seeded culture dishes, grown to 80% confluency, and then treated with stated concentrations and times of ACAT1 inhibitor Avasimibe (AV) and/or ADAM10/17 inhibitor GW280264X (Ai). Aβ_42_ was added directly to the media of treated cells at either 300 nM (BV2) or 150 nM (iMGL) and incubated for three hours (BV2) or 1.5 hours (iMGL). When stated 40 nM sTREM2 recombinant protein was added directly to the media of treated cells immediately before Aβ_42_. When stated 250 nM RAP was added to directly to the media of treated cells 30 minutes prior to the addition of Aβ_42._ For conditioned media experiments BV2 cells were sub-cultured into culture dishes and allowed to attach overnight. At 80% confluency cells were with treated with either DMSO (CNT) or 10 µM AV for 24 hours. After 24 hours media was removed and placed on BV2 cells which were simultaneously treated with either DMSO or 10 µM AV for 24 hours. 300 nM Aβ_42_ was added directly to the conditioned media of treated cells incubated for three hours. When stated 250 nM RAP was added directly to the conditioned media of treated cells 30 minutes prior to the addition of Aβ_42_.

### Cell Harvest and Preparation for Analysis

Following treatments cells were washed twice with ice-cold Dulbecco’s phosphate-buffered saline (DPBS) then lysed in radioimmunoprecipitation assay buffer (RIPA; 10 mM sodium phosphate, 150 nM sodium chloride, 2 mM EDTA, 50 nM sodium fluoride, 1% Triton X-100, 1% sodium deoxycholate, 0.1% SDS, pH 7.2) for 20 minutes to dissociate cells for protein analysis; or treated with hexane: isopropanol (3:2 v/v) to extract lipids. Lysed samples were centrifuged at 16,000xg for 20 minutes at 4°C to remove cell debris. Hexane: isopropanol was transferred to a glass vial and allowed to evaporate leaving behind the organic lipid layer.

### Cholesterol Analysis

Cholesterol levels were measured using Amplex™ Red Cholesterol Kit from Invitrogen. From the hexane: isopropanol extraction the dried organic layer was reconstituted in the provided 1x reaction buffer containing 0.1% NP-40. Samples were plated on 96-well plates then exposed to either total cholesterol reaction buffer (300 µM Amplex Red reagent, 2 U/mL HRP, 2 U/mL cholesterol oxidase, 0.2 U/mL cholesterol esterase in 1x Reaction Buffer) or free cholesterol reaction buffer (300 µM Amplex Red reagent, 2 U/mL HRP, 2 U/mL cholesterol oxidase in 1x Reaction Buffer). Fluorescence was measured using a plate reader at 530:590 nm excitation: emission. A standard curve was prepared using the standard solution included. Total and free cholesterol levels were calculated using the standard curve.

Cholesterol ester levels were calculated by subtracting free cholesterol levels from total cholesterol levels after being normalized to total protein levels from BCA analysis.

### ELISA assays

Levels of Aβ_42_ peptide in cell lysates were determined using Fujifilm Human/Rat β Amyloid (1-42) ELISA Kit Wako, high sensitivity. In brief, samples and a standard curve were plated in the antibody-coated microplate and diluted using the standard diluent provided. The plate was sealed and incubated overnight at 4°C. The solution was then discarded, and the wells were washed five times with the provided wash solution. The HRP-conjugated antibody solution was added to wells and incubated for one hour at 4°C. The antibody solution was then removed, and wells were washed five times with the provided wash solution. TMB solution was added to wells to begin the HRP reaction. Plates were incubated with TMB solution for 30 minutes at room temperature in the dark, then the stop solution was added to terminate the reaction. Absorbance was read at 450 nm using a BioTek multi-mode reader and β Amyloid (1-42) concentration were calculated using the standard curve and normalized to total protein levels from BCA analysis.

Levels of extracellular TREM2 were measured using Abcam Mouse TREM2 SimpleStep ELISA® Kit – Extracellular (Ab309115). In brief, Samples and a standard curve were prepared using the standard diluent provided as directed, media was diluted 1:100 then plated in the coated microplate. The freshly prepared antibody cocktail was added to the well and the plate was sealed and incubated one hour at room temperature on a plate shaker at 400 rpm. The solution was then discarded, and the wells were washed three times with the provided wash solution. TMB development solution was added to wells and incubated on the plate shaker for 5 minutes. Stop solution was added to terminate the reaction and absorbance was read at 450 nm. Extracellular TREM2 concentration were calculated using the standard curve and normalized to total protein levels from BCA analysis.

### Immunoblotting

Cell lysate samples were loaded and separated by 4-12% gradient Bis/Tris gels (life Technologies) and transferred to PVDF membranes using iBlot2 Transfer Stack and iBlot 2 Gel Transfer Device (Invitrogen). Blots were then incubated with SuperBlock Blocking Buffer (Thermo Fisher Scientific) for 1 hour at room temperature with shaking. Blots were then incubated with primary antibody (1:1000) overnight at 4°C with gentle shaking. Blots were washed five times before incubation with appropriate HRP-conjugated secondary antibody at room temperature for 1 hour with shaking. After 5 washes, Pierce ELC Western blotting Substrate (Thermo Scientific) was applied. Images were captured by Odyssey Platform (LI-COR) and analyzed by FIJI-ImageJ software (National Institutes of Health). If necessary, blots were then incubated with mild stripping buffer (1.5% glycine, 1% SDS, and 1% tween 20 pH 2.2) for 1 hour with shaking before re-probing with another primary antibody.

## Materials

Human/Rat β Amyloid (42) ELISA Kit Wako from Fujifilm (290-62601), Amplex™ Red Cholesterol Assay Kit (A12216), NuPAGE 4-12% Bis-Tris Gel from and iBlot2 PVDF regular stacks and anti-rabbit HRP (A16110) from Invitrogen. Albumin Standard (23210), Pierce ECL western blotting substrate, Pierce BCA Protein Assay, Pierce Premium-Grade Sulfo-NHS-SS-Biotin (PG82077) and SuperBlock™ T20 (TBS) Blocking Buffer from Thermo Scientific. Anti-TREM2 clone 78, (rat monoclonal) MABN755 and anti-rat HRP (AP136P) from EMD Millipore. Rabbit anti-beta actin (4967S) and rabbit anti-Src (2108S), rabbit anti-pSrc (6943T), rabbit anti-Syk (13198S), and rabbit anti-pSyk (920S) from cell signaling. Rabbit anti- SOAT1/ACAT1 polyclonal antibody from Cayman chemical. Full-Range protein ladder from Prometheus Genesee Scientific, Avasimibe (HY-13215) from MedChemExpress, Amyloid β-Protein (1-42) from Bachem, recombinant mouse TREM2 (19-171) His Tag (TR2-M52H3) from Acro biosystems, and RAP human recombinant (553506) EMD Millipore. Complete Mini, EDTA-Free protease inhibitor tablets (11836170001) from Roche, Dulbecco’s Modification of Eagle’s Medium (DMEM).

### Quantification and statistical analysis

The data in each figure represent biologically independent replicates. No statistical methods were used to predetermine sample size. Statistical analysis performed using GraphPad Prism (GraphPad Software). Data were analyzed by one-way ANOVA or two-way ANOVA followed by Tukey’s multiple comparisons tests or unpaired two-tailed Students t test. All p values and statistical test are indicated in the figure legend.

### Resource Availability

Request for further information and resources should be directed to and will be fulfilled by the lead contact. Plasmids and cell lines generated in this study are available from the lead contact with a completed materials transfer agreement and may require payment. All data reported in this paper will be shared by the lead contact upon request.

